# Fetal QRS Detection by Multiple Channel Temporal Pattern Search Applied to the NInFEA Database

**DOI:** 10.1101/2022.08.29.505738

**Authors:** Bruce Hopenfeld

## Abstract

**Background:** In some cases, the fetal electrocardiogram (ECG) has a relatively low signal to noise ratio. Much of the literature pertaining to fetal ECG processing focuses on methods for separating the fetal ECG from the maternal ECG rather than detecting the fetal QRS in high noise conditions. This paper describes the application of a previously described pattern search methodology (TEPS) for detecting the fetal QRS in noisy ECGs.

**Algorithm Summary:** Signals are processed in non-overlapping 5 s segments. The maternal QRS peaks are detected in a segment from a reliable lead. Next, for each of a potentially large number of leads, after removing the maternal QRS peaks from the set of searchable peaks, single channel TEPS is applied to search for provisional fetal peak sequences. The provisional sequences are scored according to previously described single channel TEPS quality measures: temporal regularity, peak pair amplitude ratios, and number of skips. The provisional sequences across all leads are aggregated according to their average RR intervals and scores. The most likely average RR interval for the segment is estimated from this aggregation by forming a score weighted average of the RR intervals. Sequences with RR intervals close to the chosen RR interval are selected to form a set of high quality sequences. To account for the possibility of peak time offsets between different channels, these sequences are time aligned by time lagged cross correlation. An optimal sequence is formed from these time aligned sequences, with optimality criteria based on peak timing coherence, temporal regularity, and skips.

**Data:** The above procedure was applied to 60 ECG signals (avg. duration approximately 31 s) from the Non-Invasive Multimodal Foetal ECG-Doppler Dataset for Antenatal Cardiology Research (“NInFEA”). The NInFEA dataset also includes ultrasound recordings from which RR intervals may be extracted.

**Results:** Out of 59 records, RR intervals were well tracked in 57 records. (One of the 60 NInFEA records was excluded due to the inability to obtain a sinus rhythm peak sequence from the ultrasound recording). For these 57 records, 84% and 95% of the ECG RR intervals were within 5 ms and 10 ms, respectively, of the ultrasound derived RR intervals. The mean and median average absolute value RR interval difference (between ECG and ultrasound) over 5 second segments were 3.2 ms and 2.4 ms respectively, with 93% of segments having a mean average absolute value RR interval difference less than 7ms.

**Conclusion:** The TEPS methodology shows promise for ECG based fetal sinus rhythm monitoring.

## 1. Introduction

Processing of the non-invasively recorded fetal electrocardiogram (ECG) remains a subject of active research [1]. Most of the literature focuses on methods for separating the fetal ECG from the maternal ECG, whether by removing the latter through adaptive filtering or by blind source separation [1]. When the fetal ECG is relatively clean, these techniques can produce very good results. However, these separation/isolation methods do not explicitly address the problem of detecting heart beats in high noise conditions, which sometimes occur in the context of fetal ECGs. For example, in Figure 1, which shows ECG signals recorded from a mother’s abdomen, removing the maternal signal or somehow isolating the “fetal signal” (assuming that’s possible) leaves open the issue of detecting the fetal QRS amongst relatively large noise peaks. Approximately 15%-25% of the records in the Non-Invasive Multimodal Foetal ECG-Doppler Dataset for Antenatal Cardiology Research (NInFEA) [2,3,4], which served as the test dataset in the present work, are characterized by a relatively low amplitude fetal ECG across a variety of abdominal leads.

**Figure 1.**
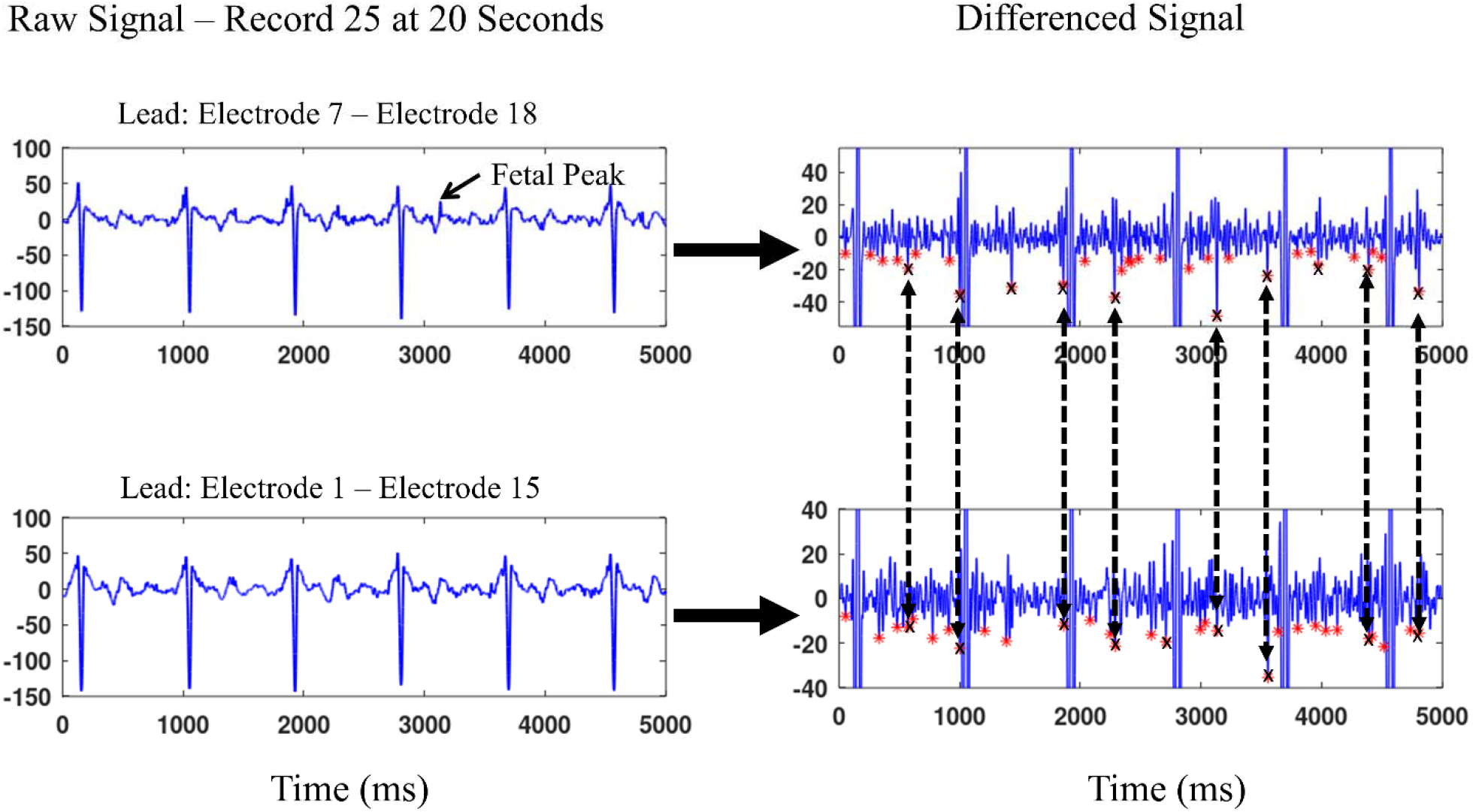
The signals in the right panels are differenced versions of the left panel (bandpass filtered) signals from record 25 of the NInFEA database. An arrow points to one of the few noticeable fetal QRS complexes visible in the non-differenced signals. In the differenced signals, the red asterisks indicate the 30 “best” peaks after the maternal peaks have been removed from consideration. Single channel TEPS was run separately on the two channels, resulting in single channel high quality peak sequences indicated by ‘X’s’. The dashed lines show the temporal correspondence between peaks in these two sequences, which in turn increases the likelihood that the sequences represent the fetal heart beat. (In the right hand panels, the maternal peaks extend beyond the y axis limits of the plots.)

A multi-channel Kalman filter [5, 6] has been applied to noisy fetal QRS detection. The state space model comprises QRS shape and RR interval dynamics. However, as pointed out by Nathan and Jafari [7], the assumptions underlying the Kalman filter, including the extended version thereof, may not be satisfied in the context of ECG signals. More particularly, in high noise conditions, many QRS complexes may be so distorted that all that remains is a simple spike, which cannot be distinguished from a noise spike by shape criteria. In the case of serial processing such as that implemented by a Kalman filter, a single false detection can throw off subsequent detections. In such conditions, schemes such as particle filtering [7] or single channel temporal pattern search (TEPS) [8, 9] that can search for combinations of peaks in multi-second long segments of the ECG will tend to outperform serial detection schemes.

Where multiple ECG channels are available, sequence score criteria should optimally combine information from all of the channels. A multi-channel sequence search approach (MTEPS) has been applied to noisy multi-channel capacitive electrodes embedded in car seats [10]. In that case, there were only three channels and the QRS peak time offsets between channels, an important component of MTEPS, was assumed to be known *a priori*. In contrast, in the NInFEA, there are a very large number of possible bipolar leads, and fetal QRS peak time alignment between leads has to be determined on the fly. The present work describes the extension of MTEPS to address these issues.

## 2. Algorithm

Figure 2 is a high-level block diagram of the algorithm, which processes data in non-overlapping five second segments. First, maternal QRS complexes are detected from a reliable lead. Next, single channel TEPS [9] is applied to each of a possibly large number of leads to generate a set of provisional peak sequences. As previously described, single channel TEPS involves selecting a set of “best” peaks (candidate peaks, as indicated by red asterisks, Figure 1 right hand panels), and searching through these candidate peaks for temporally regular sequences. (The temporal regularity constraint can be removed to detect various irregular rhythms.) Specifically, the single channel sequence score SC is set equal to [9]:

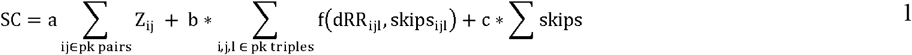

where Z is the prominence ratio for each peak pair in a sequence, dRR_ijl_ is the change in RR intervals between 3 consecutive peaks with corresponding skipped beats skips_ijl_, f() is a Gaussian with a standard deviation that depends on skips_ijl_. The values for the coefficients a, b, and c were carried over from previous work [9]:

**Figure 2.**
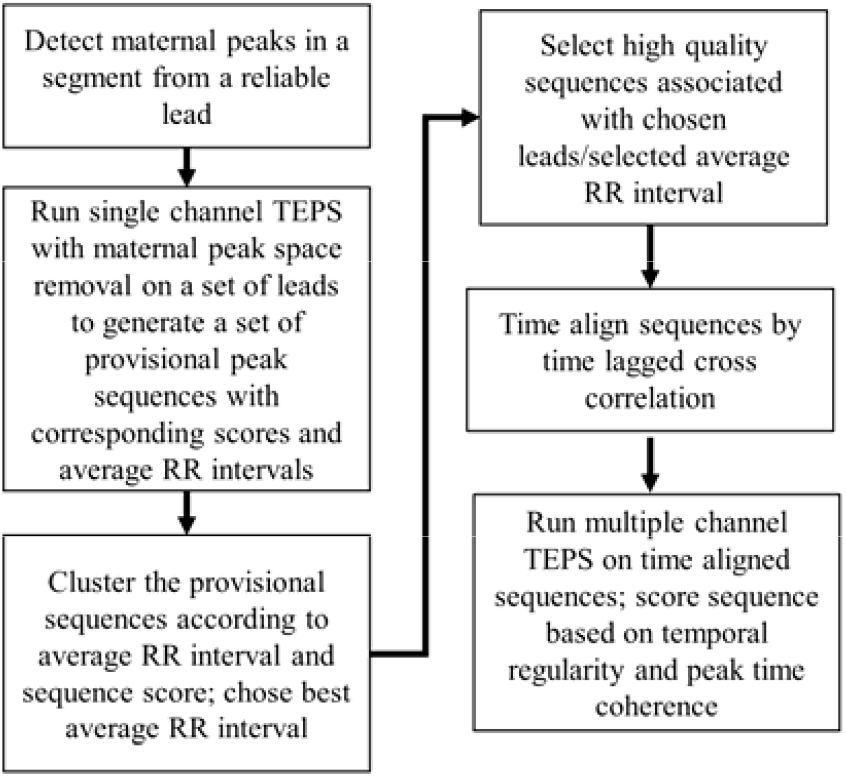
High level flowchart of multiple channel TEPS with lead selection.

The maternal peaks are effectively removed from the single channel TEPS processing: they cannot belong to the set of candidate peaks and are excluded from peak prominence (Z) scores. To exclude the maternal peaks, the reference maternal peak times are time aligned with the local channel peaks, and peaks within the latter that are close to the time aligned maternal peaks are removed. Let the reference maternal peak times be TR={tr1, tr2 … trN}, and the local channel peak times be TL-{tl1, tl2,…tlN}. (Candidate peaks are selected from TL.) An offset t_o_ is determined such that the TR peaks temporally align with the most peaks in TL. That is, t_o_ is chosen so that the number of peaks that satisfy |TR-t_o_ -TL| < th is maximized, where th is time alignment threshold. These peaks are then removed from TL. A practical scheme for carrying out this time alignment will be described below.

The above processing results, for each lead, in a set of provisional sequences characterized by average segment RR intervals and scores. For example, the sequences indicated by X’s in Figure 1 are provisional sequences with associated average segment RR intervals of approximately 430 ms. All sequences with scores above a threshold are aggregated as shown by example in Figure 3, a plot of sequence scores versus average RR intervals for the provisional sequences. (For illustrative purposes, the Figure 3 plot shows only one sequence for each lead, but multiple sequences per lead are possible.) The most likely average segment RR interval is selected by forming an average segment RR interval weighted by scores. In the right hand panel of Figure 3, a large number of sequences cluster tightly around a specific RR interval, and many of these sequences have high quality scores. In the left hand panel, corresponding to a much noisier segment, clustering is less visually obvious.

**Figure 3.**
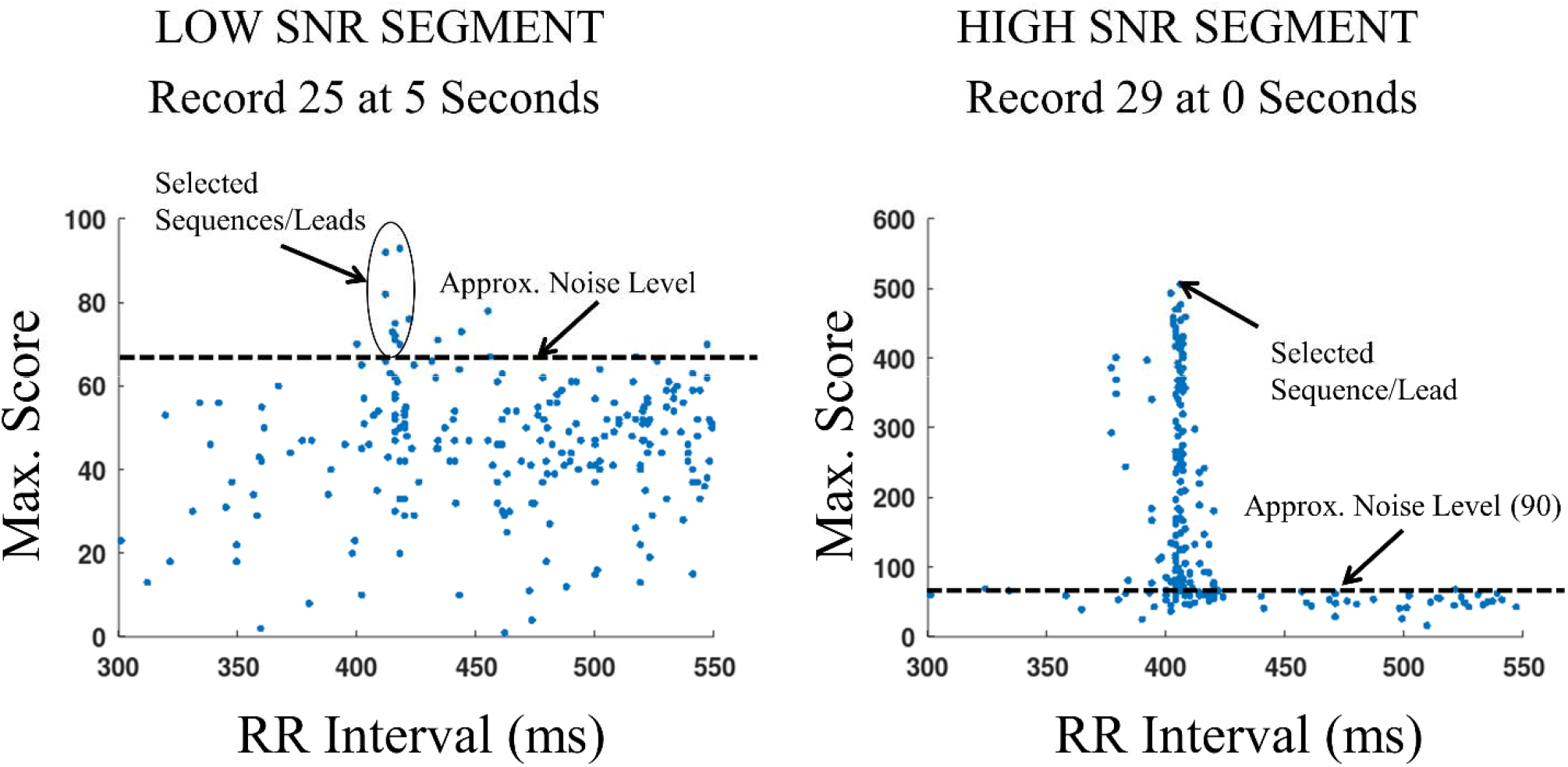
Lead selection in the case of a low SNR segment (left panel) and a high SNR segment right panel. Each dot corresponds to the best (highest scoring) sequence associated with a particular lead. Each best sequence has an associated RR interval. Leads are selected by clustering RR intervals, weighted by scores.

The high quality sequences (each corresponding to a single lead) chosen by score weighted clustering are then time aligned by performing time lagged cross correlation, and a final sequence is derived from the time aligned sequences, as will be described below. The final sequence is chosen according to temporal regularity and skips, and in addition according to peak time coherence. As described in [10], the more peaks (across channels) that occur at approximately the same time, the more likely that a true peak occurred at that time (assuming that noise is uncorrelated across channels).

The high quality sequences are time aligned by time lagged cross correlation of peak indicator signals that are generated from the sequences. Figure 4 shows an example of this technique. Three high quality sequences (corresponding to three different channels) to be time aligned are shown on the left. For each sequence, a peak indicator signal is formed with triangles centered on the peak times, as shown in the right of the Figure. The width of the triangles encodes the score for matches as a function of peak time difference. (A non-triangular shape, such as a Gaussian, could encode non-linear peak time coherence scoring.) The indicator signals are cross correlated. In Octave/Matlab, a large number of signals can be efficiently, simultaneously cross correlated by the “xcorr” function. The optimal set of time lags (offsets) would appear to be a difficult optimization problem. Instead, the lead/sequence/peak indicator signal with the maximum sum of cross correlation values is first chosen and added to the signal with the highest cross correlation. This signal is then cross-correlated with the remaining signals. The above process is repeated until all peak indicator signals have been added.

**Figure 4.**
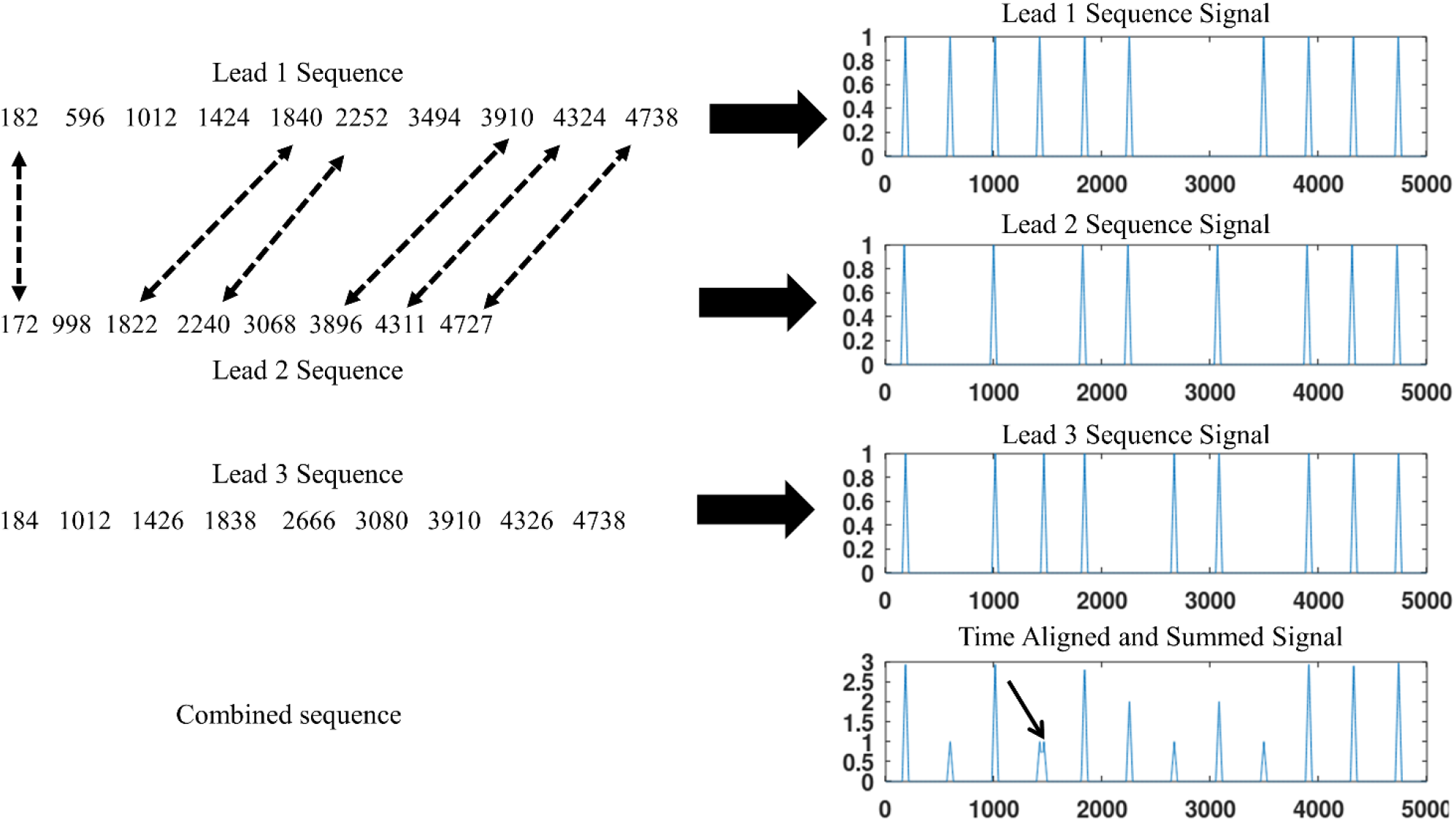
Time alignment and peak time coherence scoring of sequences by cross correlation of peak time indicator signals. The dashed arrows in the left panel show corresponding peaks in the first two sequences; the first sequences is roughly 13 ms ahead of the peaks in the second. The arrow in the lower right panel points to two possible peak locations around 1500 ms.

The result, as shown in the bottom of the right side of Figure 4, is a summed indicator signal with peak values that occur at times corresponding to averaged peak times. Further, the peak values of the summed indicator signal correspond to the peak time coherence quality score. A search is performed on the average peak times to generate sequences, which are again scored according to temporal regular and skips, and also peak time coherence. (In the example shown in Figure 4, 11 of the peaks in the summed signal clearly correspond to the optimal sequence, but the two peaks pointed to by the arrow correspond to 2 different peak times at approximately t=1500ms. The search process thus generates two different sequences corresponding to these two different peak times with all other 11 peaks in common.)

## 3. Experimental Data/Processing Details

The NInFEA dataset includes 60 records, between 7.5 s and 120 s long, corresponding to 39 pregnant women, between the 21st and 27st week of gestation [2,3,4]. Many different fetal positions are represented. “Unipolar” signals were recorded from (a) 22 electrodes on and near the abdomen; and (b) additional superior and posterior electrodes. In the present work, bipolar leads were derived from pairs of the 22 abdominal electrodes for a total of 231 possible bipolar leads.

The NInFEA dataset also includes pulsed wave Doppler (PWD) recordings, from which the ground truth RR intervals were obtained. Specifically, the upper and lower PWD envelopes were extracted with the provided (https://www.physionet.org/content/ninfea/1.0.0/code/envelope_extraction.m) Matlab/Octave scripts. Single channel TEPS was run on both the upper and lower envelope signals (with an effective sampling rate of 284 Hz). For each segment, the ground truth peak sequence was selected from the upper or lower signals according to which of the two had the highest scoring sequence (based on the Equation 1 score).

The ultrasound peak sequences were time aligned with the chosen fetal ECG sequence according to the time alignment procedure described above. In the case of skipped beats, in both the ultrasound and ECG sequences, interpolation/extrapolation was performed to ensure 1:1 alignment between the ground truth and fetal ECG sequences.

All computations were performed on a 2017 Hewlett Packard Laptop with an Intel Core i3-8130U CPU, base frequency 2.20GHz, with 8GB of RAM. The number of candidate peaks (NCP) was set equal to 30. Signals were downsampled to 512 Hz and low pass filtered to 90 Hz. The differencing parameters were carried over from [9], adjusted linearly according to the sampling rate, 512 Hz vs. 256 Hz in the prior work. All processing was done in Octave.

The noise threshold THn was chosen heuristically with regard to approximately 10 randomly selected segments. The maximum/minimum lag for cross-correlation was 20ms.

The maternal QRS complexes were recorded from “unipolar” electrode number 22 for every record.

## 4. Results

Out of 59 records, RR intervals were well tracked in 57 records. (One of the 60 records, record 9, was excluded due to the inability to obtain a ground truth sinus rhythm peak sequence.) For these 57 records, 84% and 95% of the ECG RR intervals were within 5 ms and 10 ms, respectively, of the ultrasound RR intervals. The left hand panel in Figure 5 is a histogram of the mean average absolute value RR interval discrepancy for each of 327 5 s long segments. The mean and median of this measure over 5 second segments were 3.2 ms and 2.4 ms respectively, with 93% of segments having a mean average absolute value RR interval discrepancy less than 7ms. Figure 5 also shows the root mean square difference per segment. The mean and medians of this measure over segments were 5.0 ms and 2.9 ms respectively.

**Figure 5.**
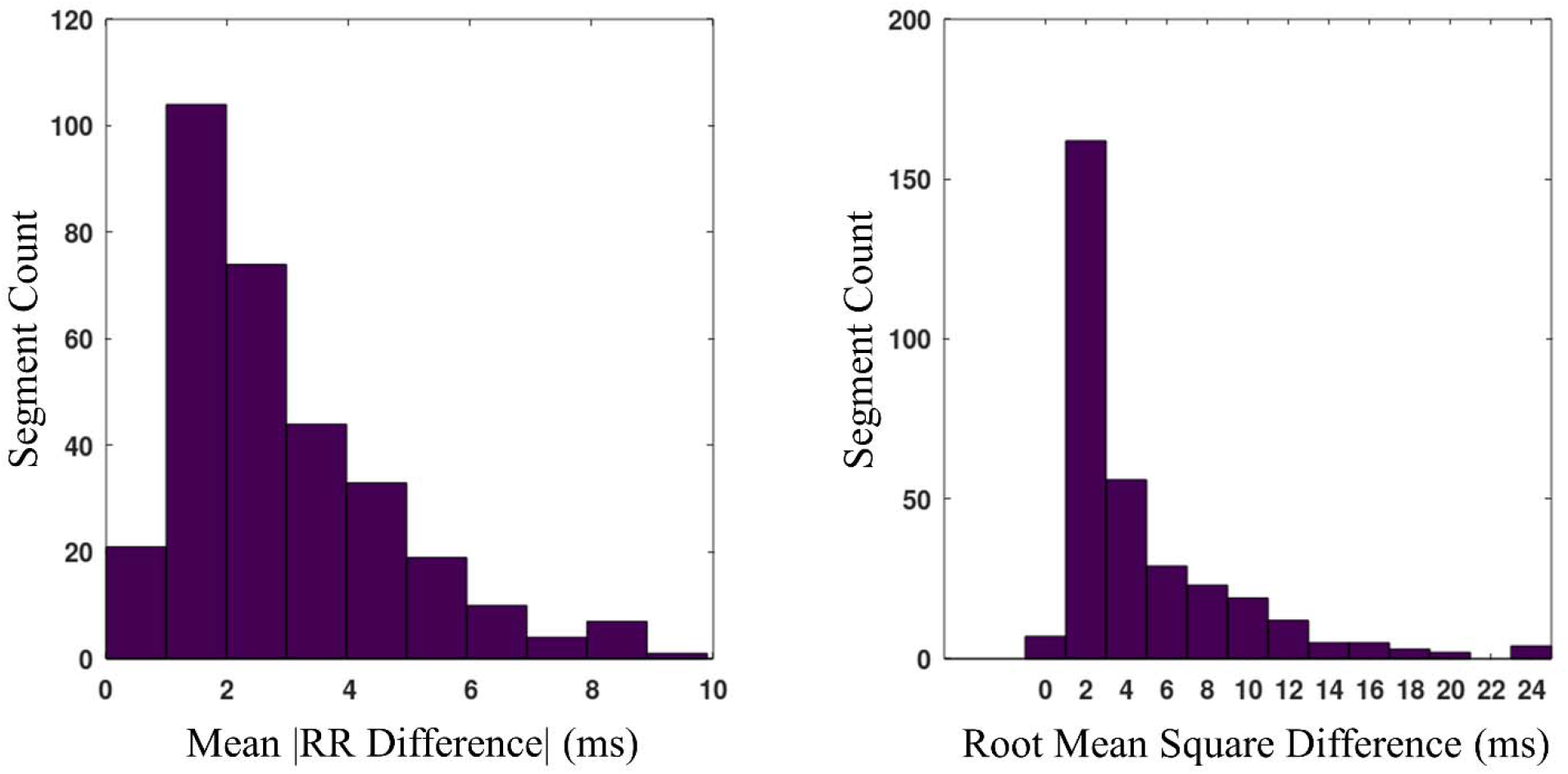
The left panel is a histogram of mean average RR difference between TEPS/ECG and ultrasound for all segments in 57 NInFEA records. The right panel is a histogram of root mean square difference over segments.

No good sequences could be found for records 12 and 18. In particular, for all segments within these records, no lead resulted in any sequence with a score that was at least as large as the noise threshold shown in Figure 3.

Appendix 1 lists all of the leads that were selected for each record. In the case of a record that required multiple leads, in most instances different subsets of the leads were chosen for different segments in the record. For 39 records, only a single lead was required, and in most of these cases, many other single leads would have produced good results. Records 21, 22, 25 and 59 required a large number of leads.

## 5. Discussion

The results suggest that fetal sinus rhythm can be accurately estimated by ECG, recorded from a supine mother, across a wide range of gestation periods and fetal positions provided that electrode coverage is sufficient. RR tracking failed in two (records 12 and18) records, each of which one of which corresponded to a high risk pregnancy with the fetus in a breach position, left, sacral anterior and right, sacral, transverse respectively. For these records, bipolar leads were formed from all combinations of the 22 abdominal electrodes and 3 posterior electrodes (numbered 23, 24, 27). It is not clear whether additional lateral and/or posterior electrodes, or more exhaustive processing, could have helped in these cases.

The ultrasound associated with record 9 was indicative of an arrhythmia, although this was not confirmed by a cardiologist. While the version of MTEPS described in this work was not configured to detect arrhythmias, the temporal regularity score may be modified or eliminated, in which case accurate detection of peaks will be based solely on peak pair prominence and peak time coherence. In this regard, by allowing RR intervals greater than the fetal maximum (approximately 550 ms), MTEPS generated sequences at an 800-850 ms period for record 9. On examination, the peaks in this record appeared to be consistent with a bigeminal type rhythm (roughly 500ms/330ms), but this could not be verified and is considered speculative, especially since the maternal rhythm was approximately 800ms.

Removing the sinus rhythm constraint to detect arrhythmias will generally increase the computational load required to find time aligned peaks but this method could be implemented by cross correlating the N “best” peaks or sequences (scored solely by peak prominence) from each lead rather than selecting only sinus rhythm sequences. If there are a large number (e.g. 100s) of leads, they could be hierarchically cross correlated in groups of leads less likely to be correlated and chosen from leads more likely to be high quality (e.g. by referring to a list like Appendix 1), given the orientation of the fetus. A zero offset could initially be applied to all leads, with a wider match window, and after each iteration, only high quality peaks would be selected for further comparison.

As described in [10], correlation of noise between channels can confound the peak time coherence analysis. In the present work, good results were obtained without analyzing noise correlation, even though many lead sets (Appedix 1) included shared electrodes, which likely resulted in substantial noise correlation. In other environmental or algorithmic circumstances, such as where the sinus rhythm constraint is removed, it may be necessary to correct for noise correlation.

Maternal QRS complexes were reliably detected from a single electrode (#22) in every record in the NInFEA dataset. If a high quality maternal rhythm ca not be detected from a single lead, the multi-channel TEPS procedure may be applied to detect the maternal heart beat sequence.

With regard to other work on the NInFEA dataset, Sulas et al.[3] reported a correlation coefficient of 0.89 between their QRS detection algorithm and ultrasound for instantaneous fetal heart rate over all records. In the present work, for the 57 records successfully processed (excluding the last few seconds of a record that did not fall into a 5 s segment), the correlation coefficient for instantaneous heart rate/RR interval was 0.99. Sulas et al.[3] derived the ultrasound RR intervals through visual inspection whereas, in the present work, that process was performed by algorithm. Keenan et al.[11] applied a prior fetal ECG extraction approach [12] to NInFEA to assess reliability of electrode/lead selection; results were reported in terms of a reliability measure based on a +/-10 beats/minute difference between ECG detected peaks. +/-10 beats/minute corresponds to an RR interval range far greater than the RR interval differences described in the present work, making a comparison very difficult. Shaham et al. [13] applied a source separation method to the NInFEA dataset but did not provide detail QRS or RR interval statistics.

## APPENDIX A SELECTED LEADS

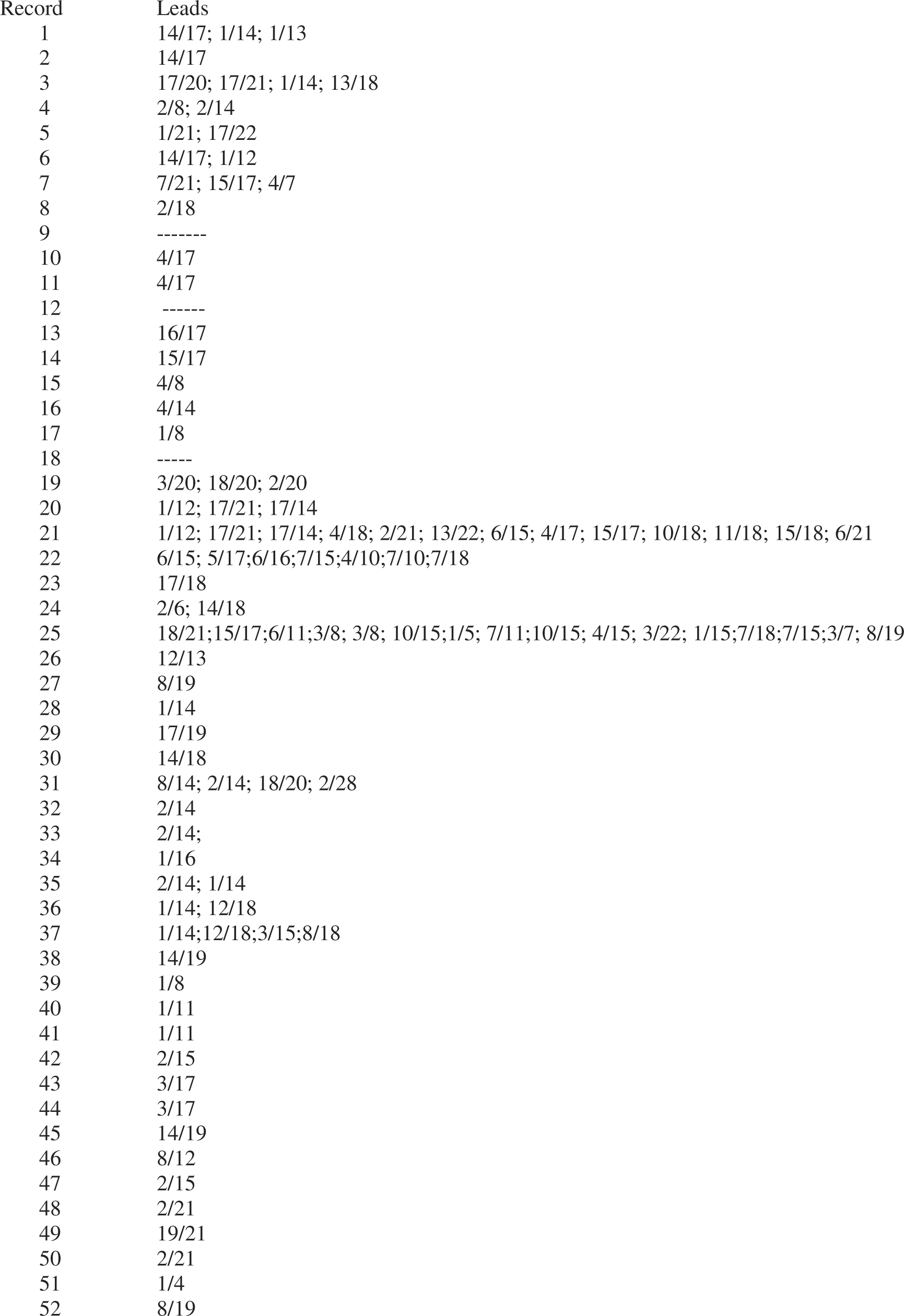
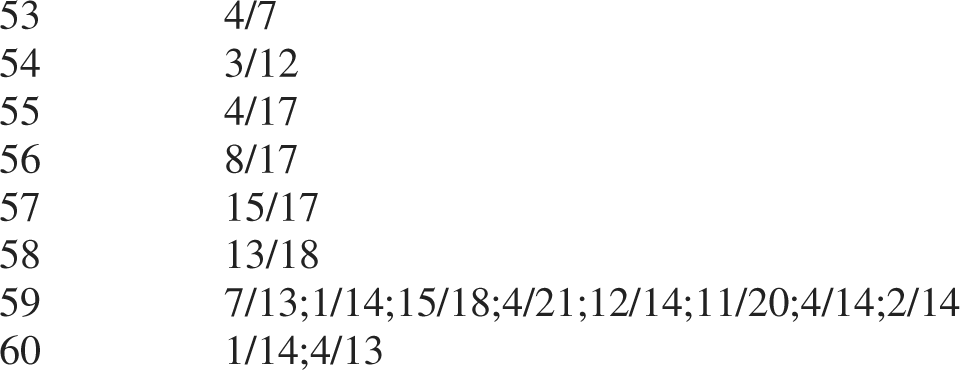

## Notes

### Competing Interest Statement

The authors have declared no competing interest.

